# *Faecalibacterium prausnitzii* induces an anti-inflammatory response and a metabolic reprogramming in human monocytes

**DOI:** 10.1101/2024.10.06.616495

**Authors:** Camille Danne, Laura Creusot, Rodrigo de Oliveira Formiga, Florian Marquet, Delphine Sedda, Laura Hua, Pauline Ruffié, Hang-Phuong Pham, Iria Alonso Salgueiro, Loic Brot, Marie-Laure Michel, Philippe Langella, Jérémie H. Lefevre, Harry Sokol, Nathalie Rolhion

## Abstract

**Background and aims:** *Faecalibacterium prausnitzii*, a highly abundant bacterium in the human gut microbiota, has been linked to overall health and is decreased in several pathological conditions, such as Inflammatory Bowel Disease (IBD). *F. prausnitzii* has shown anti-inflammatory properties in human and mouse models, notably through the induction of IL-10 signaling. Here, we investigated which cell types from human blood and intestinal tissue are responsible for producing IL-10 induced by *F. prausnitzii*, and providing the first mechanistic insights.

**Methods:** Immune cells isolated from human blood and intestinal lamina propria of patients with IBD and non-inflamed controls, were stimulated with either *F. prausnitzii* EXL01 strain or *Escherichia coli* lipopolysaccharide (LPS) and analysed by Legendplex, ELISA, flow cytometry, RNA-sequencing (RNAseq), and Seahorse technology.

**Results:** *F. prausnitzii* EXL01 strain induced the direct and dose-dependent production of IL-10 in CD14^+^ monocytes from the systemic circulation and intestinal tissue of IBD patients and non-inflamed controls, without inducing a pro-inflammatory response as compared to LPS stimulation. RNAseq analysis corroborated these results and revealed that *F. prausnitzii* EXL01 strain differentially affects cell energy metabolism compared to LPS. The anti-inflammatory response induced by *F. prausnitzii* in monocytes was dependent on mitochondrial respiration.

**Conclusion:** *F. prausnitzii* induces an anti-inflammatory response and rewires energy metabolism in human monocytes, which might explain its beneficial impact on intestinal inflammation and human health in general. These results provide new insight into the mechanisms underlying the anti-inflammatory effects of *F. prausnitzii* and are crucial for a better understanding of its potential use in the treatment of IBD.

## Introduction

Inflammatory bowel disease (IBD), including Crohn’s disease (CD) and ulcerative colitis (UC), are characterized by chronic and relapsing inflammation of the gastrointestinal tract, whose incidence and prevalence rates are increasing in both developed and developing countries^1^. There is a significant need to identify markers that could predict relapses and complications, and develop new treatments, which would have considerable socioeconomic benefits. The exact pathogenesis of IBD remains enigmatic, notably due to its multifactorial etiology. IBD partly results from genetic susceptibilities that are largely responsible for dysregulated immune responses toward commensal bacteria^2–6^. Additionally, environmental factors and lifestyle habits play a role, and the specific impact of gut microbiota alterations on IBD pathogenesis has been highlighted in numerous studies^7,8^. Indeed, the gut microbiota appears as both a marker of inflammation and an actor in IBD development. The pro-inflammatory effect of IBD-associated microbiota has been demonstrated by microbiota transfer from humans to mice^9^ and by the positive effects of fecal microbiota transplantation used as a treatment of IBD in clinical trials^10,11^. One of the most significant and consistent findings regarding the alterations of the gut microbiota in IBD is the decreased abundance of *Faecalibacterium prausnitzii*, a member of the Firmicutes phylum, belonging to the Clostridium IV group^12–14^. *F. prausnitzii* is the most abundant bacterial species in the human intestinal microbiota of healthy adults, representing around 5% of the total fecal microbiota^15–18^. Its high prevalence and abundance indicate its significant contribution to microbiota functions in healthy individuals. Indeed, high levels of *F. prausnitzii* are associated with health, whereas low levels are linked to diseases such as IBD^17,19^ and cancer^20,21^. Moreover, this bacterium has shown immunomodulatory effects both *in vitro* and *in vivo* in different colitis mouse models^17,22–24^. The anti-inflammatory effect of *F. prausnitzii* notably relies on the induction of the production^1^ of the cytokine IL-10 by immune cells, together with the low induction of pro-inflammatory cytokines^17,25^. *F*. *prausnitzii* promotes tolerogenic IL-10- producing dendritic cells that in turn favour the priming of IL-10–producing CD4^+^ T cells *in vitro* in human^26^. Moreover, *F*. *prausnitzii* induces IL-10-producing CD4^+^CD8α^+^ human Tregs that protect against intestinal inflammation in a humanized murine model^19,27,28^. However, the complete picture of the cell types responding to *F. prausnitzii* by producing IL-10 in humans, and the mechanisms at play, remain unknown.

This work used *F. prausnitzii* EXL01 strain, already in clinical development for IBD (NCT05542355). We started with an unbiased approach to investigate which cell types from human blood respond to *F. prausnitzii* EXL01 strain by inducing IL-10 together with a low pro-inflammatory response. We found that CD14^+^ monocytes are the main producers of IL-10 in response to *F. prausnitzii* EXL01 strain, with a lower production of IL-23 and TNF-α compared to lipopolysaccharide (LPS). We confirmed the anti- inflammatory response induced by *F. prausnitzii* EXL01 strain in CD14^+^ monocytes isolated from the ileum and colon lamina propria of IBD patients and non-inflamed controls. To explore the mechanisms involved, we performed RNA sequencing (RNAseq) on CD14^+^ monocytes isolated from both human blood and intestinal tissue from IBD patients. We found that *F. prausnitzii* EXL01 strain does not induce the pro-inflammatory response triggered by the classical bacterial factor LPS. In addition, *F. prausnitzii* EXL01 strain acts on cell energy metabolism of human CD14^+^ monocytes from both peripheral blood and ileal tissue, inducing oxidative phosphorylation and inhibiting glycolysis and several cell death-related pathways, including apoptosis, necrosis and senescence, when compared to LPS stimulation. Using Seahorse technology, we confirmed that the anti-inflammatory response to *F. prausnitzii* EXL01 strain relies on mitochondria and the rewiring of cellular energy metabolism in human CD14+ monocytes. This study provides new insights into the mechanism underlying the anti-inflammatory effects of *F. prausnitzii* on human immune cells.

## Results

### *Faecalibacterium prausnitzii* EXL01 strain induces the production of IL-10 by human CD14 ^+^ monocytes

To investigate the immunomodulatory properties of *F. prausnitzii* in humans, peripheral blood mononuclear cells (PBMCs) were isolated from the fresh blood of six healthy volunteers and stimulated for 16 h with *F. prausnitzii* EXL01 strain at different multiplicities of infection (MOI 1, 10 and 100; corresponding to Fp1, Fp10, Fp100 in figures) or with *Escherichia coli* K12 lipopolysaccharide (LPS, 100 ng/ml), used as a positive control (**Fig. 1**). Release of cytokines in cell supernatant using an unbiased approach showed that *F. prausnitzii* EXL01 strain induced the secretion of the anti-inflammatory cytokine IL-10 in a dose-dependent manner, whereas the induction of the pro-inflammatory cytokines IL-23 or TNF-α notably was reduced compared to the LPS condition, leading to increased IL-10/IL-23 and IL-10/TNF-α anti-inflammatory ratios (**Fig. 1A**). ELISA analyses performed independently confirmed this result (**Suppl. Fig. 1A**). To identify the specific cell types involved in this immune response, stimulated PBMCs were analyzed by flow cytometry after IL-10 intracellular staining (**Suppl. Fig 1B**). *F. prausnitzii* EXL01 strain induced a dose-dependent increase of the percentage of IL-10^+^ cells in CD14^+^ monocytes, and to a lesser extent in CD4^+^CD8^+^ double positive T cells (DP8) (**Fig. 1B**). No significant increase in IL-10^+^ cells was observed for CD4^-^CD8^-^ double negative (DNeg), CD4^+^ and CD8^+^ T cells, and CD19^+^ B cells (**Fig. 1B**).

**Figure 1.**
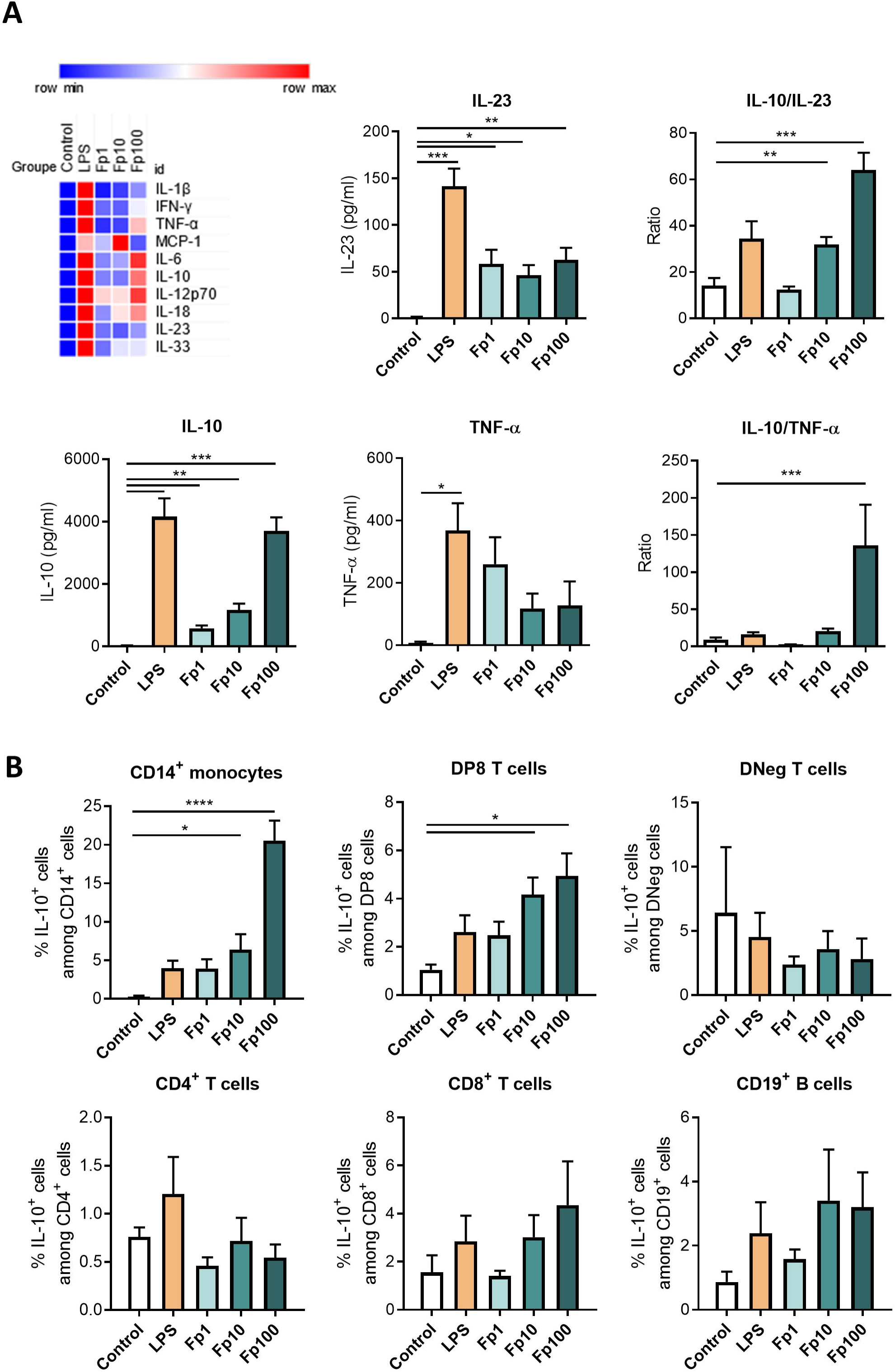
*Faecalibacterium prausnitzii* EXL01 strain induces the production of IL-10 in PBMCs from systemic circulation. (A) Peripheral blood mononuclear cells (PBMCs) from the fresh blood of healthy volunteers were isolated and stimulated for 16 h with *F. prausnitzii* EXL01 strain at different multiplicities of infection (MOI 1, 10 and 100; corresponding to Fp1, Fp10 and Fp100) or with *Escherichia coli* K12 lipopolysaccharide (LPS, 100 ng/ml), used as a positive control. (Upper left panel) Heatmap representing Legendplex analysis (N=6), blue referring to minimal induction, red to maximal induction. (Other panels) Cytokine concentrations and anti-inflammatory ratios for IL-10, IL-23 and TNF-α. (B) Flow cytometry analysis of PBMCs stimulated for 16 h in the same conditions as in (A), showing IL-10^+^ cells among different immune cell populations. N=5. DP8, double positive CD4^+^CD8^+^ T cell population; DNeg, double negative CD4^-^CD8^-^ T cell population. Data are mean ± SEM of two independent experiments. *P<0.05, **P<0.01, ***P<0.001 and ****P<0.0001, as determined by Mixed-effects analysis and Dunnett’s multiple comparisons test (A) and ordinary one-way ANOVA (B).

### *F. prausnitzii* EXL01 strain directly induces the production of IL-10 in CD14 ^+^ **monocytes from systemic circulation.**

Intracellular staining was performed for the three cytokines IL-10, IL-23 and TNF-α together with an antibody panel differentiating classical (CD14^+^CD16^-^), intermediate (CD14^+^CD16^+^) and non-classical (CD14^-^CD16^+^) monocyte populations (**Suppl. Fig. 2A**). As observed with LPS, the stimulation of PBMCs with *F. prausnitzii* EXL01 strain increased the percentage of classical and non-classical monocytes, and significantly reduced the percentage of intermediate monocytes, leading to a slight decrease in the percentage of total CD14^+^ monocytes (including classical and intermediate populations, **Suppl. Fig. 2B-C**). *F. prausnitzii* EXL01 strain induced an increase in IL-10^+^CD14^+^ monocytes, and a slight increase of IL-23^+^CD14^+^ monocytes, but no increase in TNF-α^+^CD14^+^ monocytes, leading to a dose-dependent but non statistically significant increase in both IL-10/IL-23 and IL-10/TNF-α anti-inflammatory ratios (**Fig. 2A**). Similar trends are obtained for the three different monocytes subtypes taken individually (data not shown).

**Figure 2.**
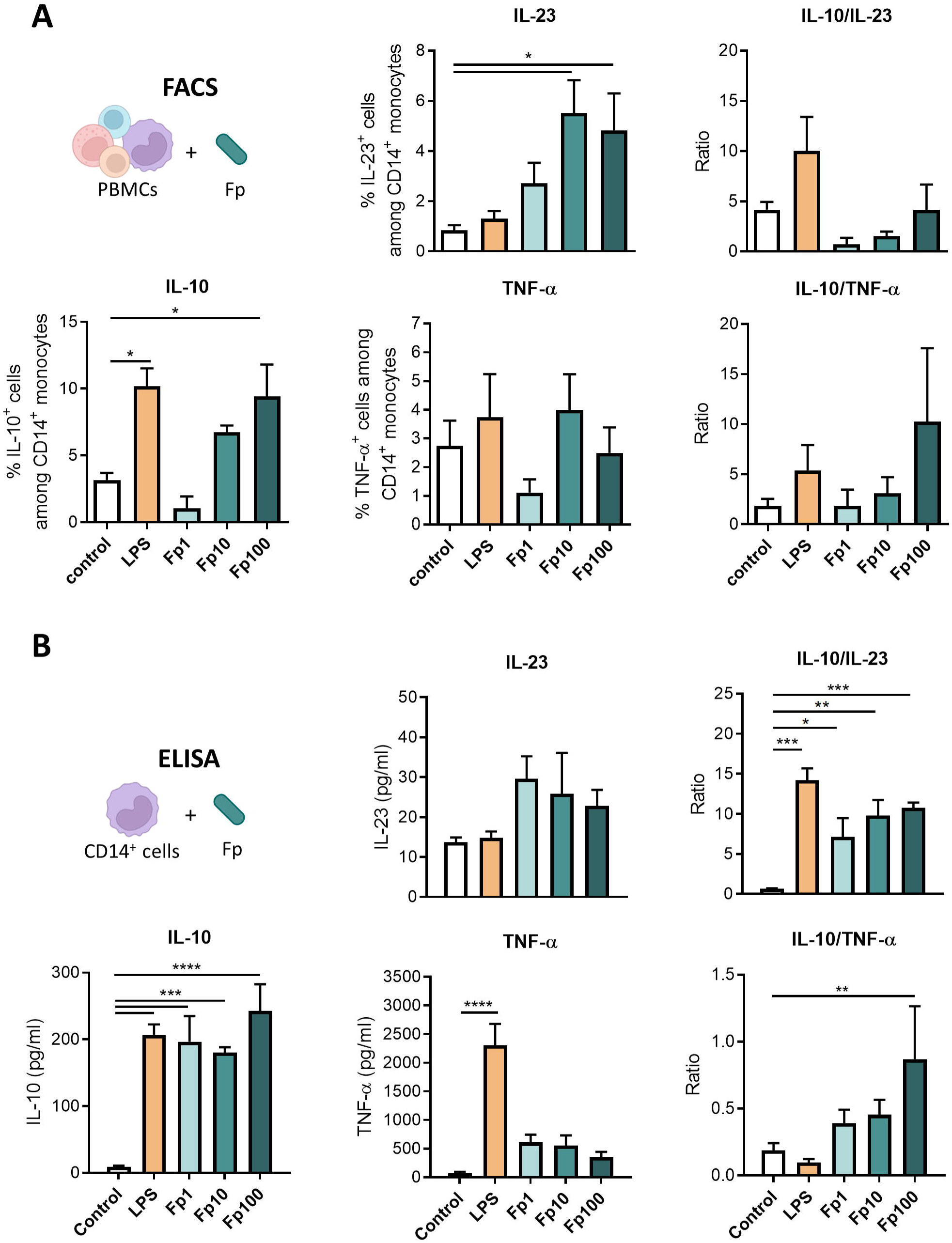
*F. prausnitzii* EXL01 strain directly induces the production of IL-10 in CD14+ monocytes from systemic circulation. (A) Flow cytometry analysis of PBMCs stimulated in the same conditions as in Fig.1, showing IL-10^+^, IL-23^+^ and TNF-α^+^ cells among CD14+ monocytes and anti-inflammatory ratios (N=4). (B) Cytokine concentrations and anti-inflammatory ratios for IL-10, IL-23 and TNF-α obtained by ELISA analysis (corrected by LDH) of the supernatant of CD14+ monocytes isolated from PBMCs and stimulated for 16 h in the same conditions as in Fig.1 and 2A. Data are mean ± SEM of two independent experiments. *P<0.05, **P<0.01, ***P<0.001 and ****P<0.0001, as determined by two-way ANOVA (A) and ordinary one-way ANOVA (B).

Moreover, the stimulation of CD14^+^ monocytes purified from total human PBMCs recapitulated the capacity of *F. prausnitzii* EXL01 strain to induce IL-10 secretion by these cells, together with low production of IL-23 and TNF-α compared to LPS, leading to a dose-dependent increase in anti-inflammatory ratios (**Fig. 2B, Suppl. Fig. 2D**). Interestingly, the lower dose of *F. prausnitzii* EXL01 strain Fp1 was sufficient to induce maximal production of IL-10 in CD14^+^ monocytes, and increasing doses reduced the induction of pro-inflammatory cytokines in a dose-dependent manner (**Fig. 2B**). Thus, the production of IL-10 in CD14^+^ monocytes relies on a direct stimulation by *F. prausnitzii* and does not depend on other immune cell types (**Fig. 2B**).

### *F. prausnitzii* induces the production of IL-10 in CD14 ^+^ **monocytes from intestinal tissue of patients with IBD and non-inflamed controls**

To evaluate the immunomodulatory properties of *F. prausnitzii* EXL01 strain directly in the human intestinal tract, including in pathological conditions, we took advantage of surgical intestinal tissues, both ileum and colon, from patients operated for IBD or colon adenocarcinoma (ADK). We used inflamed but non-ulcerated intestinal tissue from patients with IBD, and tumor-free healthy margins of ADK patients as non-inflamed controls. Intestinal mucosa containing epithelial and lamina propria immune cells were isolated, and stimulated 0.5 mm^2^ pieces with three doses of *F. prausnitzii* EXL01 strain (as used in our cellular *in vitro* assay: Fp1, Fp10, Fp100), or with LPS for 16 h (**Fig. 3**). Despite a complex tissue structure and a large cellular diversity, *F. prausnitzii* EXL01 strain induced IL-10 secretion both in total ileum and colon mucosa (**Fig. 3B**), and similarly in IBD and ADK patients (**Suppl. Fig. 3**). In comparison, the pro-inflammatory cytokines IL-23 and TNF-α were only slightly induced in response to *F. prausnitzii* EXL01 strain in this model, leading to dose-dependent increases in anti-inflammatory ratios (**Fig. 3C**, significant only for IL-10/IL-23 in ileum).

**Figure 3.**
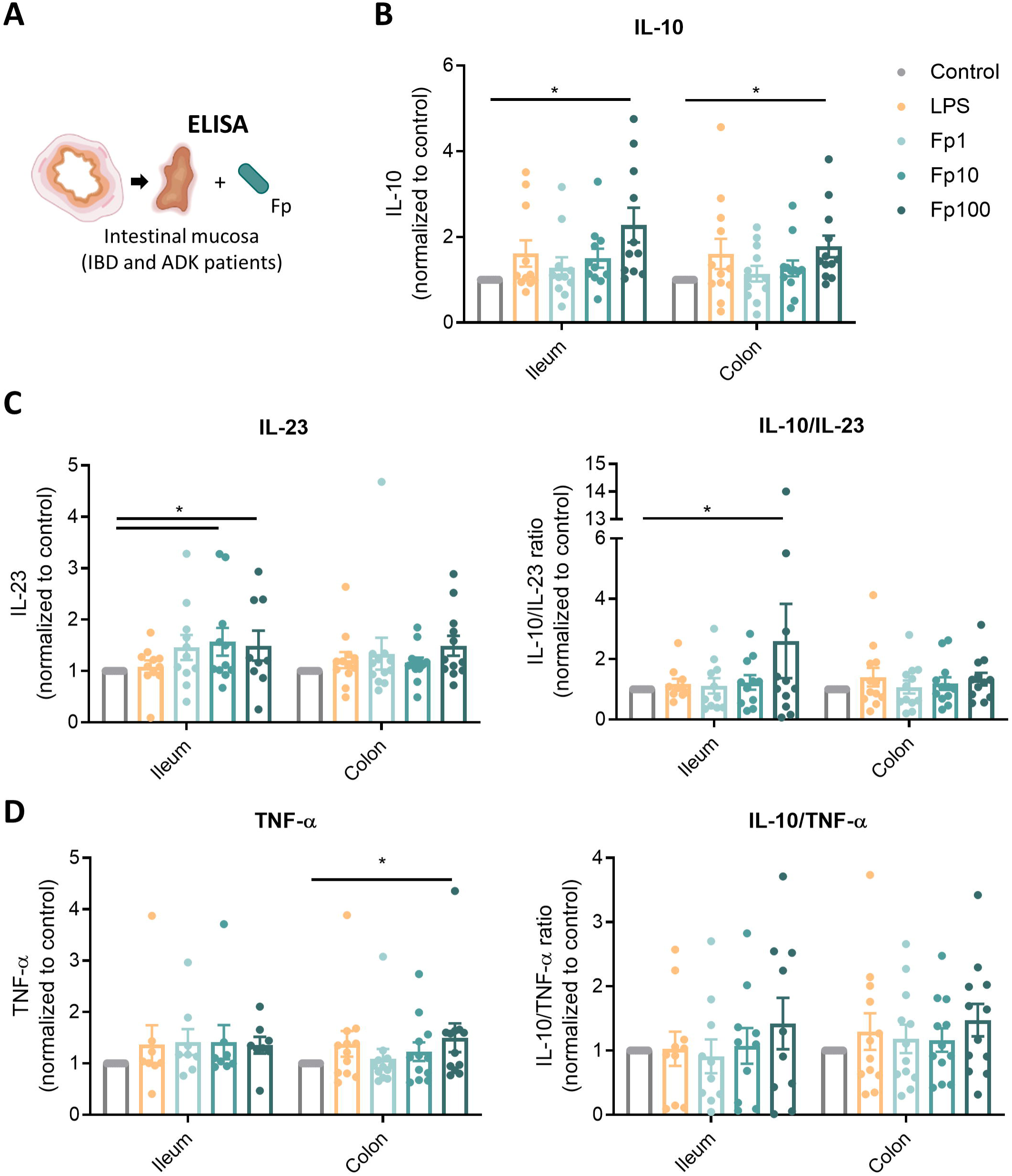
*F. prausnitzii* induces the production of IL-10 in intestinal tissue of IBD patients and non-inflamed controls. (A) Schematic representation of the experiment setup. IL-10 (B), IL-23 (C) and TNF-α (D) concentrations and corresponding anti-inflammatory ratios measured by ELISA (corrected by LDH) in the supernatant of intestinal mucosa (ileum and colon) from IBD and non-inflamed controls (adenoma-carcinoma or ADK patients), stimulated for 16 h with different doses of *F. prausnitzii* EXL01 strain (Fp1, Fp10, Fp100) or LPS, as performed previously. N=23 (IBD ileum N=6; IBD colon N=8; ADK ileum N=5; ADK colon N=4). Data are mean ± SEM. *P<0.05, as determined by two-way ANOVA.

To directly assess the response of intestinal immune cells to *F. prausnitzii* EXL01 strain, immune cells were isolated from fresh surgical tissue lamina propria, and were stimulated in similar conditions (**Fig. 4**). *F. prausnitzii* EXL01 strain induced an increase in IL-10^+^CD14^+^ monocytes in a dose-dependent manner mostly in ileum cells, and similarly in IBD and ADK patients (**Fig. 4B, Suppl. Fig. 4A-B**). *F. prausnitzii* EXL01 strain induced a decrease in IL-23^+^CD14^+^ or TNF-α^+^CD14^+^ monocytes especially in ileum, leading to increased anti-inflammatory ratios in ileum (**Fig. 4C-D**). As observed with PBMCs (**Fig. 1B**), *F. prausnitzii* EXL01 strain did not induce the production of IL-10 in CD3^+^ T cells and CD19^+^ B cells isolated from intestinal tissue in our assay (**Suppl. Fig. 4C**).

**Figure 4.**
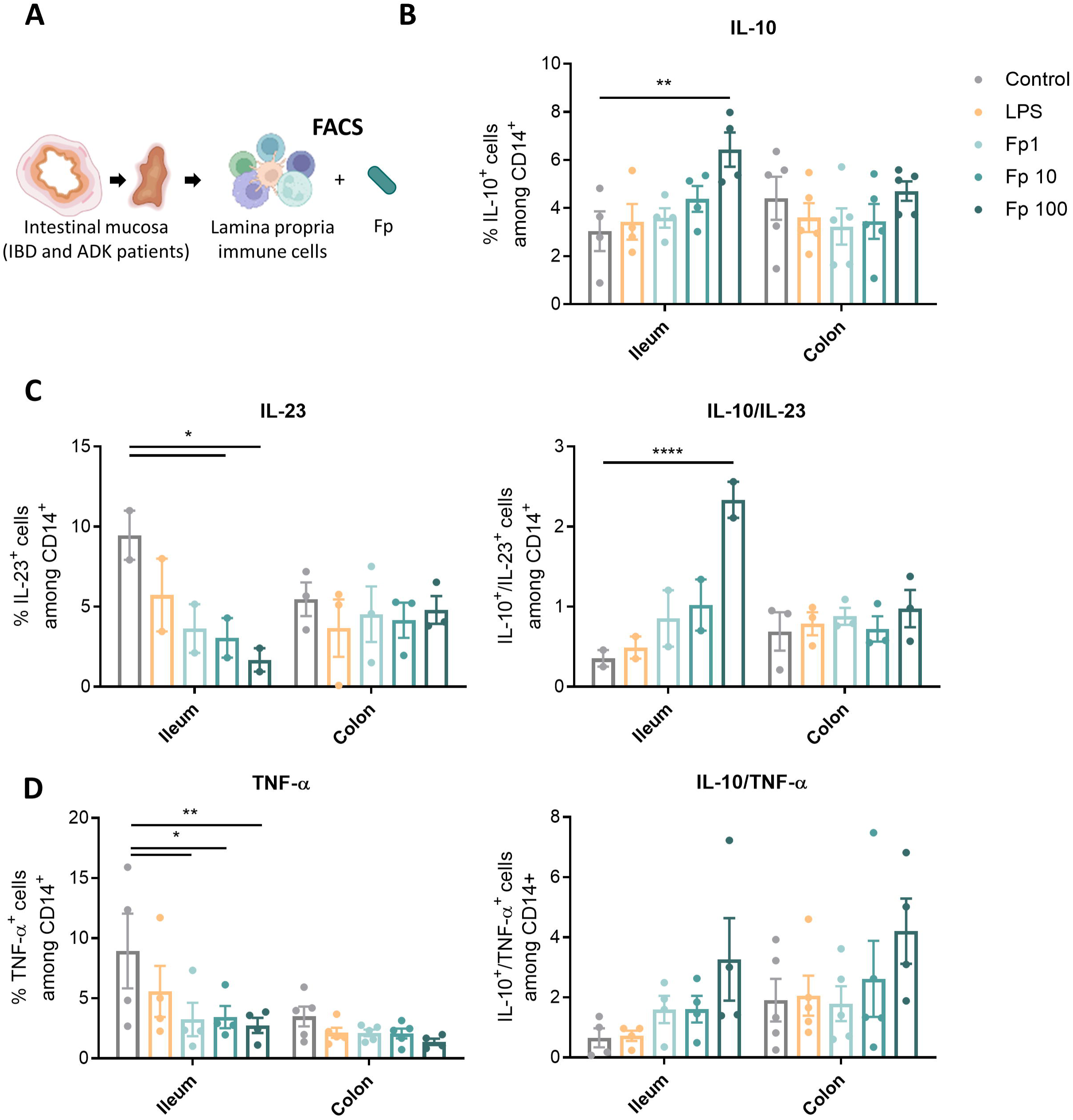
*F. prausnitzii* induces the production of IL-10 in CD14 ^+^ monocytes isolated from intestinal lamina propria of IBD patients and non-inflamed controls. (A) Schematic representation of the experiment setup. Percentage of IL-10^+^ (B), IL-23^+^ (C) and TNF-α^+^ (D) cells among CD14^+^ monocyte population measured by flow cytometry in the total lamina propria immune cells isolated from the intestinal mucosa (ileum and colon) of IBD and ADK patients, and stimulated for 16 h with different doses of *F. prausnitzii* EXL01 strain (Fp1, Fp10, Fp100) or LPS, as performed previously. N=9 (IBD ileum N=3; IBD colon N=2; ADK ileum N=1; ADK colon N=3). Data are mean ± SEM. *P<0.05, **P<0.01, and ****P<0.0001 as determined by two-way ANOVA.

Altogether, these results confirm that *F. prausnitzii* EXL01 strain triggers an anti-inflammatory response in CD14^+^ monocytes from both the systemic circulation and the human intestinal tissue.

### *F. prausnitzii* EXL01 strain differentially affects immune response and cell energy metabolism compared to LPS

To get an insight into the mechanisms underlying the anti-inflammatory effects of *F. prausnitzii* on CD14^+^ monocytes, RNA-sequencing (RNAseq) analysis was performed on CD14^+^ cells sorted from human PBMCs (**Fig. 5**, **Suppl. Fig. 5**) and stimulated with *F. prausnitzii* EXL01 strain (Fp10 and Fp100) or LPS. Principal component analysis (PCA) showed that stimulation of CD14^+^ monocytes with *F. prausnitzii* EXL01 strain induced a different gene expression profile compared to the classical bacterial factor LPS (**Fig. 5B**). Functional analysis (Over-representation analysis [ORA] of differentially expressed gene [DEG], Reactome pathway database) confirmed that both *F. prausnitzii* EXL01 strain and LPS could induce IL-10 signaling pathway (**Suppl Fig. 5A-B**). However, many pro-inflammatory pathways, such as Interferon_Gamma_Response, Interferon_Alpha_Response, Tnfa_Signaling_Via_Nfkb, Inflammatory_Response, Complement, Il6_Jak_Stat3_Signaling and IL2_Stat5_Signaling, were significantly down-regulated by *F. prausnitzii* EXL01 strain compared to LPS (ORA, DEG Reactome and Hallmark pathway databases, **Fig. 5C and Suppl Fig. 5C**). Interestingly, cell metabolism-related pathways were also differentially regulated between the two conditions, with an up-regulation of Oxydative_Phosphorylation and a down-regulation of Apoptosis in the Fp100 condition versus LPS (**Fig. 5C**). Difference-in-differences analysis supported the finding that *F. prausnitzii* EXL01 strain did not induce pro-inflammatory pathways compared to LPS, but up- regulated Oxidative Phosphorylation while down-regulating Glycolysis and Apoptosis pathways (**Fig. 5D-E**).

**Figure 5.**
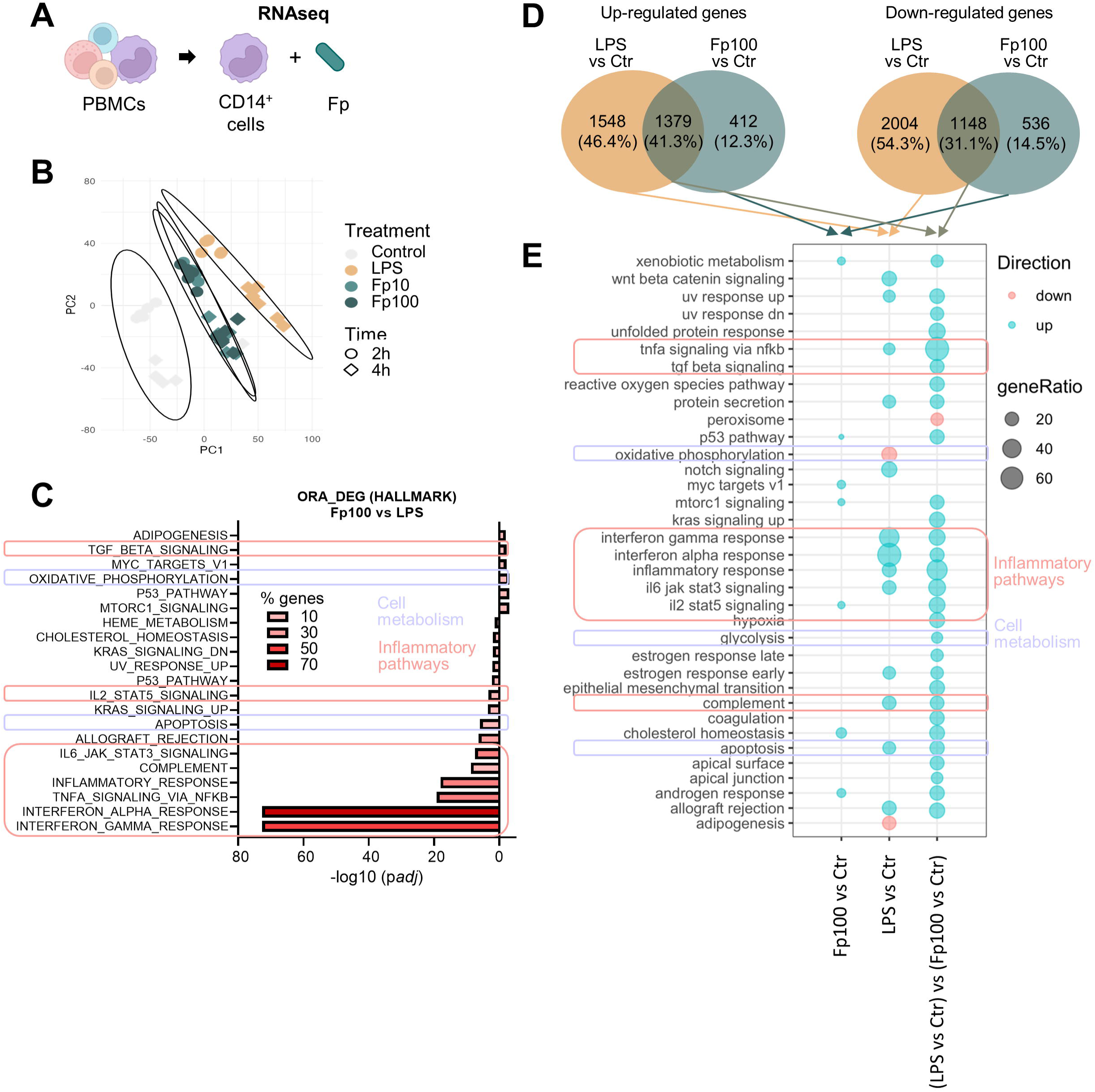
*F. prausnitzii* EXL01 strain differentially affects immune response and cell energy metabolism compared to LPS in CD14 ^+^ monocytes from peripheral human blood. (A) Schematic representation of the analysis setup. (B) Principal Component Analysis showing the repartition of the expression profiles of blood CD14^+^ monocytes according to their treatment, i.e. different doses of *F. prausnitzii* EXL01 strain (Fp10, Fp100) or LPS, as performed previously, for 2 and 4 h. (C) Over-representation analysis of differentially expressed genes (DEG) in the Fp100 compared to the LPS condition at 4 h on Hallmark pathway database. Left are pathways enriched by down-regulated genes in Fp100 compared to LPS, right pathways enriched by up-regulated genes. Percentages of the DEGs involved in each pathway are indicated by the red color intensity. (D) Venn diagrams representing up- and down-regulated genes in Fp100 versus Ctr (Control) conditions, LPS versus Ctr conditions, and between the (LPS vs Ctr) and (Fp100 vs Ctr) conditions. (E) Bubble plot showing significantly enriched pathways by up-regulated (in blue) and down-regulated (in red) genes in DEG lists. Bubble size shows the ratio of DEG genes involved in the pathways. Inflammatory pathways are encircled in pink, cell metabolism-related pathways in purple. N=6.

To go further, RNAseq analysis on CD14^+^ monocytes isolated from ileum lamina propria of patients with IBD (n=3) was performed in similar experimental conditions (**Fig. 6A**). The activation of IL-10 signaling pathway by *F. prausnitzii* EXL01 strain was confirmed (gene set enrichment analysis, Reactome database, **Fig. 6B**). Moreover, difference-in-differences analysis pointed out a down- regulation of pro-inflammatory pathways, including Interferon_Gamma_Response, Interferon_Alpha_Response, Tnfa_Signaling_Via_Nfkb, Inflammatory_Response and Il6_Jak_Stat3_Signaling in Fp100 compared to LPS, together with a down-regulation of Apoptosis (**Fig. 6C**).

**Figure 6.**
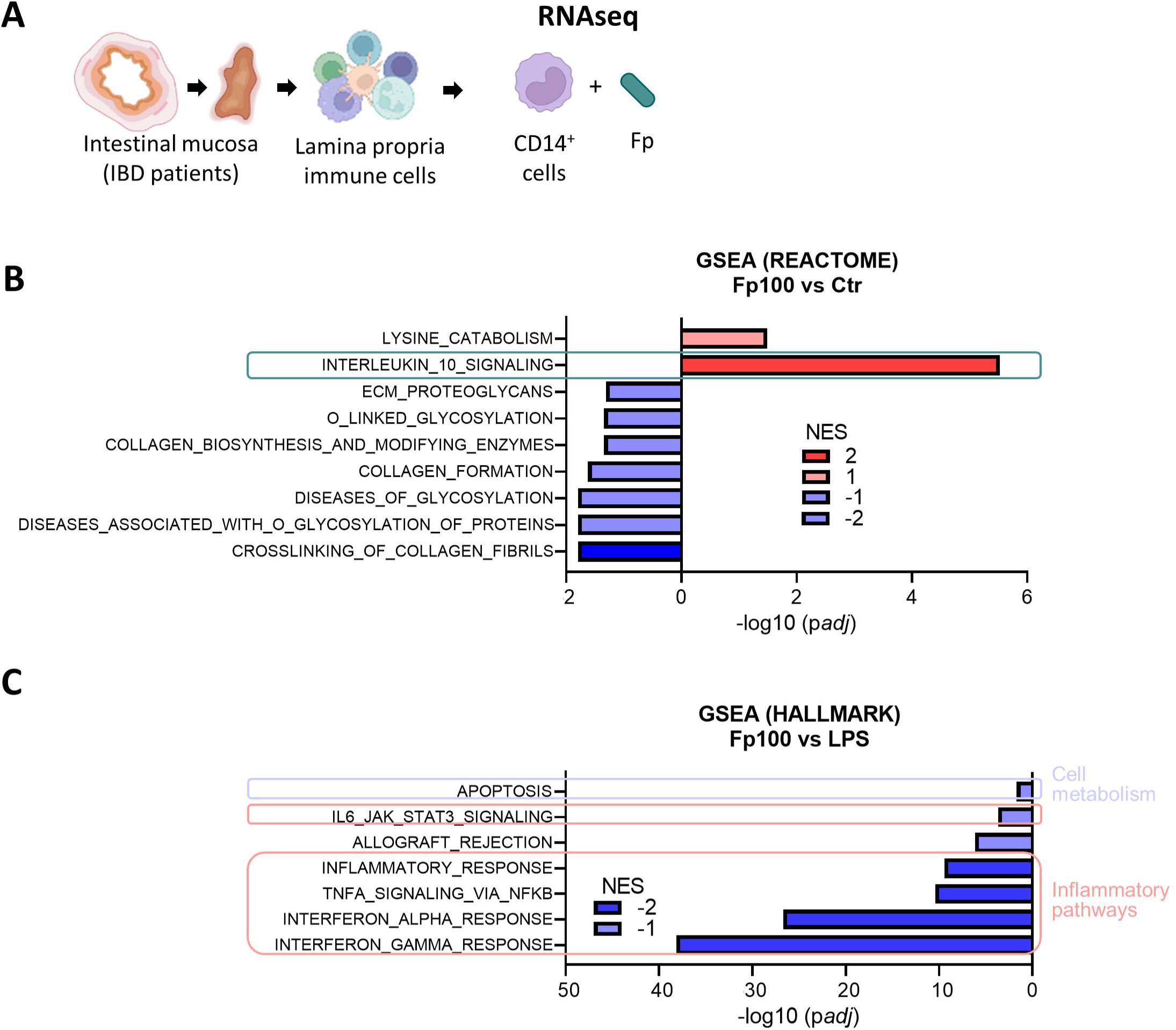
*F. prausnitzii* EXL01 strain differentially affects immune response and cell energy metabolism compared to LPS in CD14 ^+^ monocytes from ileal human tissue. (A) Schematic representation of the analysis setup. CD14^+^ monocytes were isolated form the ileal lamina propria of IBD patients, and stimulated for 4 h with different doses of *F. prausnitzii* EXL01 strain (Fp10, Fp100) or LPS, as performed previously. (B) Significant enriched REACTOM pathways in the comparison between Fp100 and Ctr conditions at 4 h identified by GSEA. Interleukin_10_Signaling pathway is encircled in green. (C) Significant enriched HALMARK pathways in the comparison between Fp100 and LPS conditions at 4 h by GSEA. Left/blue are down-regulated pathways in Fp100 compared to Ctr or LPS conditions, right/red up-regulated pathways. NES, Normalized Enrichment Score. Inflammatory pathways are encircled in pink, cell metabolism-related pathways in purple. N=3.

### Anti-inflammatory effects of *F. prausnitzii* EXL01 strain relies on rewiring energy metabolism in CD14^+^ monocytes

Guided by RNAseq results, we further explored the impact of *F. prausnitzii* EXL01 strain on the energy metabolism of CD14^+^ monocytes. Cells isolated from fresh human PBMCs were stimulated in the presence or absence of oligomycin, a complex V mitochondrial respiration inhibitor (**Fig. 7A, Suppl. Fig. 6A-C**). Interestingly, oligomycin reduced IL-10/TNF-α anti-inflammatory ratio induced by *F. prausnitzii* EXL01 strain, but not by LPS, suggesting that the anti-inflammatory response found here is dependent on mitochondrial function contrary to LPS (**Fig. 7A**). Real-time bioenergetic profile analysis using Seahorse technology showed that basal stimulation with *F. prausnitzii* EXL01 strain, particularly Fp100, increased oxidative phosphorylation (OXPHOS) assessed through the oxygen consumption rate (OCR) measurement (**Fig. 7B, Suppl. Fig. 6B**. Besides, Fp100 modulated OCR levels in response to oligomycin, FCCP and Rot/AA, showing effects on OXPHOS parameters, including basal respiration, ATP production, maximal respiration and spare capacity of CD14^+^ cells, contrary to LPS stimulation (**Fig. 7B-C, Suppl. Fig. 6B-C**). On the other hand, LPS stimulation strongly decreased the above-mentioned parameters and increased non-mitochondrial respiration (corresponding to aerobic glycolysis), as previously demonstrated^29,30^ (**Fig. 7B-C**). Non-mitochondrial respiration is typically attributed to inflammation-associated enzymes, including cyclo-oxygenases, lipoxygenases, and NADPH oxidases, and increases with reactive oxygen species^31,32^; thus, it is considered as a negative indicator of the energetic health of the cell. Reciprocally, *F. prausnitzii* EXL01 strain in basal conditions slightly decreased aerobic glycolysis in a dose-dependent manner, as measured by the extracellular acidification rate (ECAR), whereas LPS stimulation increased this parameter (**Fig. 7B**). Interestingly, increasing doses of *F. prausnitzii* EXL01 strain blocked LPS-induced glycolysis activation and OXPHOS inhibition (**Fig. 7B-C**).

**Figure 7.**
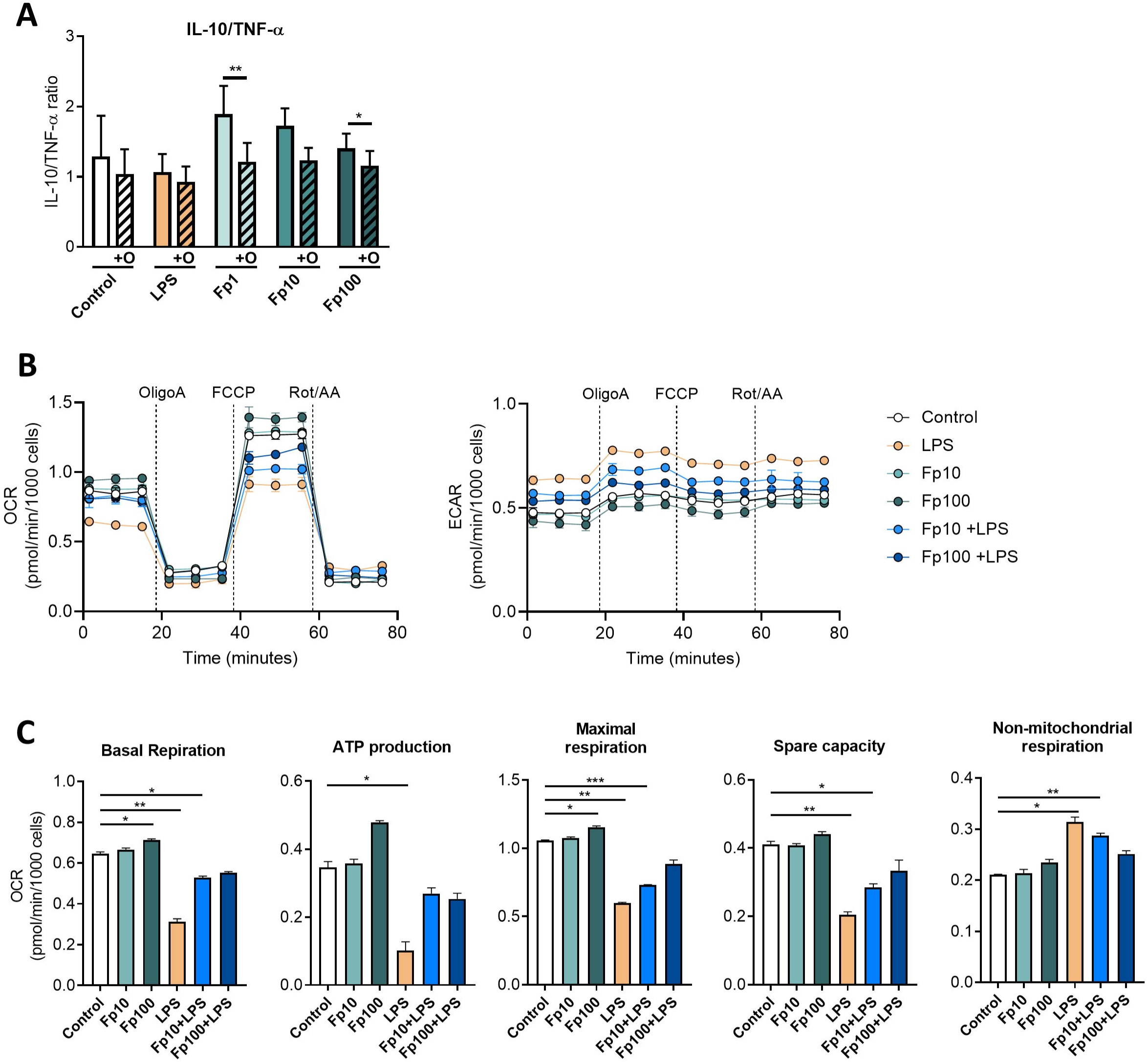
*F. prausnitzii* EXL01 strain relies on mitochondrial activity to induce an anti-inflammatory response in CD14 ^+^ monocytes. (A) IL-10/TNF-α anti-inflammatory ratio as measured by ELISA in the supernatant of CD14^+^ monocytes purified from PBMCs and stimulated for 16 h as previously, except from an additional step of 1 h pre-treatment with oligomycin, an inhibitor of oxidative phosphorylation (ATP synthase). (B) Oxygen consumption rate (OCR) and Extracellular acidification rate (ECAR) of CD14^+^ blood monocytes stimulated for 6 h with different doses of *F. prausnitzii* EXL01 strain (Fp10 and Fp100) or LPS as performed previously, or with the combination of Fp10 or Fp100 and LPS (Fp10+LPS and Fp100+LPS), measured during a Seahorse Cell Mito Stress assay. (C) Basal respiration, ATP production, maximal respiration, spare capacity and non-mitochondrial respiration obtained from the Seahorse Cell Mito Stress assay, as represented on the schematic OCR graph in Suppl. Fig. 7. N=3. *P<0.05, **P<0.01 and ***P<0.001, as determined by paired t test (A) or mixed-effects analysis and Dunnett’s multiple comparisons test (C).

Taken together, these results demonstrate that *F. prausnitzii* EXL01 strain promotes IL-10 production in monocytes without inducing the pro-inflammatory response triggered by LPS. Moreover, it is interesting to note that the functional changes in terms of immune response are associated with more profound cellular metabolic modifications, as previously described in the case of other pro or anti-inflammatory stimuli able to reprogram immune cell phenotypes and functions^33^.

## Discussion

*F. prausnitzii* is recognized as one of the most important bacteria of the human gut microbiome and its anti-inflammatory properties make it a good candidate for the treatment of IBD. The anti-inflammatory effects of *F. prausnitzii* are related to its ability to induce the production of IL-10 by immune cells, but the cell types and the mechanisms underlying these effects, particularly in human settings, remain largely unknown. In this work, we confirmed that DP8 cells from human PBMCs produce IL-10 in response to *F. prausnitzii.* We identified CD14^+^ monocytes as the primary IL-10 producers in response to *F. prausnitzii* in human PBMCs. *F. prausnitzii* EXL01 strain can induce high IL-10 production in a direct and dose-dependent manner in these cells, which is not associated with a pro-inflammatory response, as triggered by the bacterial factor LPS. This result was confirmed on CD14^+^ monocytes isolated not only from fresh human blood, but also from the lamina propria of ileum and colon mucosa of patients with IBD and non-inflamed controls right after surgery. Using inhibitor and Seahorse technology, we found out that, in parallel to immunomodulatory properties, *F. prausnitzii* also induces profound changes in the energy metabolism of monocytes, with increased mitochondrial activity, as opposed to LPS. Moreover, we showed that the anti-inflammatory response induced by *F. prausnitzii* EXL01 strain is dependent on mitochondrial activity.

Under environmental stimulation, immune cells undergo functional changes, from resting to activation state, which relies on reprogramming of their energy metabolism^33^. The main metabolic pathways involved in immune metabolism are glycolysis, the tricarboxylic acid (TCA) cycle, the pentose phosphate pathway (PPP), fatty acid oxidation (FAO) and synthesis, and amino acid (AA) metabolism. For instance, macrophage immune functions are associated with their energy metabolism. In homeostasis, macrophages mostly use TCA cycle, but their metabolism is modified when activated by the bacterial product LPS^34,35^. Known as M1 or classically activated macrophages, LPS-activated macrophages display pro-inflammatory properties. Their TCA cycle is altered, glucose uptake is enhanced and switches to glycolysis, which is associated with the activation of the PPP for biosynthesis of biomolecules and ATP production. In contrast, IL-4 activated macrophages, namely M2 or alternately activated macrophages, have anti-inflammatory and anti-parasitic effects, and play roles in wound healing^36^. Moreover, M2 or alternately activated macrophages show different metabolic characteristics compared to M1, with glycolysis used to support oxidative phosphorylation, leading to enhanced FAO and oxidative phosphorylation^36^. Our results suggest that the *F. prausnitzii* EXL01 strain reprograms host intestinal immune cells toward an alternatively activated phenotype with increased mitochondrial activity, although the precise mechanisms and dialogue involved remain to be elucidated.

The gut microbiota is a major actor in the modulation of both immunity and cellular energy metabolism. Despite the crucial role of the cellular energy metabolism in the ability to mount an appropriate immune response, the interactions between microbiota and immune cells regulation remain poorly understood. Most studies have focused on surface polysaccharides, including LPS, and in microbiota–derived molecules, either produced or transformed by microorganisms, such as short-chain fatty acids (SCFA), tryptophan metabolites, lipids and bile acids (BA)^7,37–41^.

Studies of immune cell energy metabolism revealed the metabolic mechanisms underlying disease progression, notably inflammatory diseases such as autoimmune diseases, chronic viral infections, and cancer^33^. In the case of IBD, a link has been found between mitochondrial dysfunction and disease severity, mostly in intestinal epithelial cells^37,42–46^. It is thus crucial to decipher more precisely the impact of host-microbiota interactions in health and disease, especially in terms of immune and metabolic regulation. The intrinsic diversity of the actors within the gut microbiota and the immune system brings an additional level of difficulty in the exploration of this crosstalk. However, this effort is crucial to identify innovative therapeutic targets for microbiota-associated diseases.

Thus, the *F. prausnitzii* EXL01 strain, well characterized and already in the clinical development stage for IBD (NCT05542355), is an ideal candidate strain for the development of LBP in the context of intestinal inflammation. Oral administration of a *F. prausnitzii-* based product could alleviate IBD patient symptoms and lead to more extended remission periods. In the cancer context, we also demonstrated recently that the *F. prausnitzii* EXL01 strain boosts the effect of an immune checkpoint inhibitor *in vitro* and in mouse models^21^, and it is currently being evaluated in several clinical trials (NCT06448572; NCT06551272, NCT06253611).

## Material and Methods

### Human samples

All human samples were collected with informed consent. Approval for human studies was obtained from the local ethics committee (Comité de Protection de Personnes Ile de France IV, IRB 00003835 Suivitheque study; registration number 2012/05NICB + Comité de Protection de Personnes Ile de France III, Biomhost study; EUDRACT number 2018-A02978-47 -18.1). All individuals (both IBD and ADK patients) from the Biomhost study were recruited in the Department of Digestive Surgery of the Saint Antoine Hospital (Paris, France). All samples were obtained between October 2022 and October 2023 (**Table 1**).

**Table 1.** Patients for ELISA and flow cytometry on total intestinal lamina propria (Biomhost cohort). Control: Healthy margin of patients operated on for colon adenocarcinoma. IBD treatment for ELISA experiments: 5-ASA, hydrocortisone, prednisone, tofacitinib; for flow cytometry experiments: 5-ASA, prednisone, azathioprine, adalimumab, tofacitinib. IBD, inflammatory bowel disease; CD, Crohn disease; UC, ulcerative colitis; SD, standard deviation; NA, not applicable; BMI, body mass index.

|  | ELISA on total mucosa |  | Flow cytometry on lamina propria |  |
| --- | --- | --- | --- | --- |
|  | IBD | Control | IBD | Control |
| n | 8 | 8 | 3 | 4 |
| UC/CD | 7/1 | NA | 2/1 | NA |
| Male Gender | 4 | 2 | 2 | 3 |
| Mean Age (SD) | 48,1 (16,1) | 60,5 (21,2) | 41,3 (11,2) | 65,6 (34,6) |
| Mean BMI (SD) | 25 (7) | 26,1 (8,2) | 26,9 (12) | 24,5 (3,4) |
| Patient with IBD treatment at time of surgery | 4 | NA | 2 | NA |
| Type of tissue analyzed |  |  |  |  |
| Colon | 3 | 6 | 2 | 3 |
| Ileum | 7 | 2 | 2 | 1 |

**Table 2.**
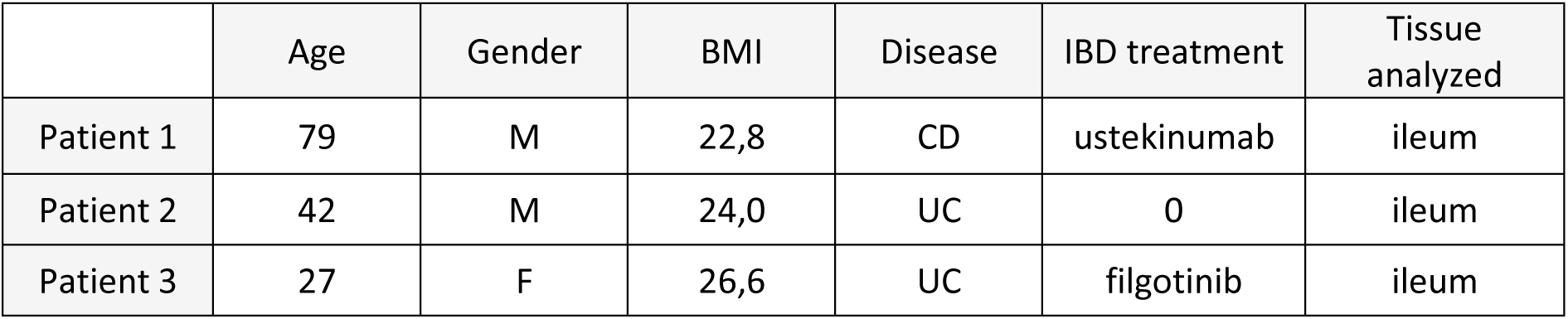
Patients for RNAseq on ileal lamina propria CD14+ cells (Biomhost cohort). M, male; F, female; BMI, body mass index; CD, Crohn disease; UC, ulcerative colitis.

|  | Age | Gender | BMI | Disease | IBD treatment | Tissue analyzed |
| --- | --- | --- | --- | --- | --- | --- |
| Patient 1 | 79 | M | 22,8 | CD | ustekinumab | ileum |
| Patient 2 | 42 | M | 24,0 | UC | 0 | ileum |
| Patient 3 | 27 | F | 26,6 | UC | filgotinib | ileum |

### Preparation of *F. prausnitzii* EXL01 culture and stimulation

*F. prausnitzii* EXL01 strain was grown at 37°C in a Exeliom Bisociences’ proprietary medium according to an in-house production process by Exeliom Biosciences. The test material was prepared to be ready-to-use, characterized by enumerating the number of morphologically intact bacteria i.e. the Total Cell Count (TCC). Total human PBMCs (5x 10^5^),), intestinal lamina propria (5x10^5^) or CD14^+^ isolated cells (1x10^5^) were treated with control medium, LPS (100 ng/ml) or EXL01 strain at 3 doses (MOI 1, 10 and 100) for 6 or 16 h as specified in the figure legend.

### Legendplex and ELISA

Legendplex (Human Inflammation Panel 1, 740809 BioLegend) and ELISA (Invitrogen, ThermoFisher Scientific) assays on cell supernatants were performed according to the manufacter recommendations. Legendplex experiments have been acquired on a BD Accuri^TM^ C6 flow cytometer and analysed using the online software Qognit. To normalized ELISA assays, Cytotoxicity Detection Kit (LDH, 11644793001, Roche) was used at the end of each experiment on 25 μl cell supernatant.

### PBMCs and CD14 ^+^ cells isolation

Fresh PBMCs were isolated from human blood using Histopaque®- 1077 (Sigma-Aldrich) and Sepmate-50^TM^ tubes (STEMCELL Technologies) according to manufacturer recommendations. PBMCs were directly stimulated or counted to isolate CD14^+^ monocytes using human CD14 MicroBeads and MS columns on a MiniMACS^TM^ Separator (Miltenyi Biotech).

#### Human lamnia propria cells isolation from intestinal resection

The intestinal mucosa was dissociated from the resection and washed for 15 min in a cocktail of antibiotics and antifungal (100 U/ml penicillin/streptomycin, 100 μg/ml gentamycin and 0.1 μg/ml amphotericin B). Fragments of 0.5 mm^2^ were cut and incubated two consecutive times in dissociation medium (PBS 1X + Hepes 10 mM + EDTA 5 mM + RPMI 2 %) for 15 min at 37 °C under 100 rpm agitation. Tissue was harvested and incubated in digestion buffer (RPMI + FCS 2% + DNaseI (0.5 mg/ml, Sigma 11284932001) + Collagenase IV (0.5 mg/ml, Sigma C5138)) for 40 min at 37°C under 100 rpm agitation. Supernatant was filtered, lamina propria immune cells counted for direct stimulation or isolation of CD14^+^ cells as described in the section above.

### Flow cytometry

Extracellular staining was performed in FACS Buffer (PBS1X + EDTA 0.5 mM + FCS 2%). Staining includes a viability dye (Zombie Aqua Fixable Viability Kit (Biolegend) and the following antibodies: CD4 (BV605, SK3, Biolegend 344646), CD8 (BV510, SK1, Biolegend 344732), CD3 (PB, OKT3, Biolegend 317314), CD19 (AF700, SJ25C1, Biolegend 363024), CD14 (PE/Dazzle^TM^594, HCD14, Biolegend 325634), CD16 (BV785, 3G8, Biolegend 302046), HLADR (APC, L243, Biolegend 307610), Live/Dead (Zombie NIR™ Fixable Viability Kit, Biolegend 423105) for the PBMCs panel, and CD14 (PE/Dazzle^TM^594, HCD14, Biolegend 325634), CD3 (APCvio770, RE1613, Miltenyi Biotec 130113136), CD19 (BV605, HIB19, Biolegend 302244), CD16 (BV785, 3G8, Biolegend 302046) for the CD14+ monocytes panels. For intracellular staining, Cytofix/Cytoperm (BD) was used following manufacturer’s protocols with following antibodies: IL-10 (PE, FES3-9D7, Biolegend 501404), IL-23 (DyLight488, Novus 727753, FAB1716k) and TNF-α (AF700, MAb11, Biolegend 502928). All data are acquired with a CytoFlex (BD) and analyzed with FlowJo software.

### section

#### RNA-sequencing analysis

RNA was extracted from 1x10^5^ CD14^+^ monocytes using Qiagen RNeasy kit and RNA-sequencing (including a Smarter stranded V3 library preparation step) has been performed at the platform Genotyping/Sequencing, ICM Paris Brain Institute (Hôpital de la Pitié-Salpêtrière CNRS UMR 7225 – Inserm U 1127 – Sorbonne Université UM75) on an Illumina Novaseq X sequencer. Illumina’s adapters and bad quality bases (Phred < 20) were removed from reads using TrimGalore v0.6.7^47^. fastQC v 0.12.0 was used for the quality control of raw and trimmed paired-end fastqs. Salmon v 1.10.1^48^ was used to quantify reads against a mapping-based index built from the GENCODE v45 (GRCh38) transcript set^49^. Quantified transcripts were imported into R using the package tximport v1.30.0^50^. Gene-level *DESeqDataSet* object was built from previously imported transcript abundances using the package DESeq2 v1.42.1^51^ in order to perform the differential expression analysis. A pre-filtering was applied to remove (i) genes having 0 count in more than 60% of samples within one of the biological conditions and (ii) genes without official symbol. For visualisation, raw counts were transformed by the “variance stabilizing transformation” method. Principal Component Analysis (PCA) was performed on variance stabilizing transformation and normalised count dataset using the function *prcomp* to explore the relationship between biological conditions. Differential expression analysis was performed by the function *DESeq* of the package DESeq2. Briefly, (i) the size factor was estimated for each sample, (ii) the dispersion was estimated for each gene, (iii) all counts were fitted to a Negative Binomial Generalized Linear Model and (iv) contrasts and Wald’s test were performed for testing the difference of expression between biological conditions. Venn’s diagrams were used to visualize the overlap between significant genes lists through the package ggvenn v0.1.10. In case of 3 samples by group, EBSeq v2.0.0^52^ was used for differential analysis as advised by Li et al^53^. Biological function and pathways enriched by significant genes were assessed using over- representation analysis (ORA) on HALLMARK and REACTOME database through the package msigdbr v7.5.1. ORA was performed separately for significantly up-regulated genes (logFC > 0) and down- regulated genes (logFC < 0) in each comparison using the function *fora* from the R package fgsea^54^. In addition, pathways were tested for their significant enrichment between biological conditions by Gene Set Enrichment Analysis (GSEA)^55^ using the function *fgsea* from the R package fgsea. Plots were produced by the package ggplot2 v3.5.1. Benjamini-Hochberg’s method was used for controlling the False-discovery rate (FDR). Significant level was fixed at type I error alpha = 0.05.

### Seahorse experiments

Mito Stress Test assay was performed on a XF96 Extracellular Flux Analyzer (Seahorse Biosciences). CD14^+^ monocytes (1×10^5^ cells/well) were isolated from fresh human PBMCs isolated from different donors and stimulated for 6 h, as previously described. Then, cells were washed and seeded in poly-D-lysine pre-coated Seahorse plates with Agilent Seahorse XF RPMI Medium pH 7.4 supplemented with XF 10 mM Glucose Solution, XF 1 mM Pyruvate Solution, and XF 2 mM Glutamine. Then, cells were washed in Seahorse RPMI medium and incubated for 1 h at 37°C without CO_2_. In the analyzer, oligomycin 1.5 μM, FCCP 1 μM and rotenone+antimycinA 0.5 μM were injected at the indicated times. Groups were set as quadruplicates and standardization was performed after each experiment using Hoescht labeling of cells and counting in a Cytation5 microscope (Agilent Biotek) with no noticeable differences in cell numbers according to the stimulation conditions.

### Biostatistics

Data were analyzed using Prism version 8 (Graphpad Software, San Diego, USA). Values are expressed as mean ± Standard error of the mean (SEM). The statistical tests used are indicated in the figure legends. For p value: *: p<0.05, **: p<0.01, ***: p<0.001, ****: p<0.0001.

## Availability of the data, analytic methods and study materials

All data related to this study will be made available to other researchers (RNA sequencing, accession number pending).

## Authors’ contributions

C.D., H.S. and N.R. designed the study, performed the analyses, interpreted the results and drafted the manuscript. C.D., L.C., R.F., F.M. performed experiments; D.S., L.H., I.A.S. and L.B. provided technical help. H.P.P. performed the RNAseq analysis. J.H.L. provided intestinal tissue from patients.

M.L.M. generated preliminary data. P.R., M.L.M. and P.L. provided intellectual input. All authors reviewed and approved the manuscript.

## Acknowledgments

The authors thank the patients recruited in the Digestive Surgery Department of Saint-Antoine Hospital (AP-HP, Paris, France) and the nurses; Sandrine Truong (Gastroenterology Department of Saint-Antoine Hospital); Pr Mohamad Mohty and Dr Béatrice Gaugler (Saint-Antoine Research Center, Paris, France) for the use of the CytoFLEX flow cytometer; Yannick Marie, Delphine Bouteiller and Emeline Mundwiller (Platform ICM Paris Brain Institute) for RNA sequencing; and Benjamin Hadida for helpful discussion.

## Supplementary figure legends

**Supplementary figure 1.** (A) Cytokine concentrations and anti-inflammatory ratios for IL-10 and IL-23 obtained by ELISA analysis (corrected by LDH) in the supernatant of CD14^+^ monocytes isolated from PBMCs and stimulated in the same conditions as in Fig.1 and 2 (N=8). *P<0.05, **P<0.01 and ***P<0.001 as determined by mixed-effects analysis and Dunnett’s multiple comparisons test. (B) Gating strategy for flow cytometry analysis of the different immune cell populations in PBMCs.

**Supplementary figure 2.** (A) Gating strategy for flow cytometry analysis of monocytes in PBMCs. (B) FACS plot representing CD14 and CD16 expression in PBMCs stimulated by different doses of *F. prausnitzii* EXL01 strain (Fp1, Fp10, Fp100) or LPS. (C) Percentage of classical (CD14^+^CD16^-^), intermediate (CD14^+^CD16^+^), non-classical (CD14^-^CD16^+^) and total CD14^+^ monocytes among live stimulated PBMCs. (D) FACS plot representing the percentage of CD14^+^ cells before and after isolation of CD14^+^ cells from PBMCs using magnetic beads.

**Supplementary figure 3.** IL-10 concentrations measured by ELISA (corrected by LDH) in the supernatant of intestinal mucosa (ileum and colon) from non-inflamed controls (adenoma-carcinoma or ADK patients) and IBD patients showed separately, stimulated for 16 h with different doses of *F. prausnitzii* EXL01 strain (Fp1, Fp10, Fp100) or LPS, as performed previously. N=23 (IBD ileum N=6; IBD colon N=8; ADK ileum N=5; ADK colon N=4). Data are mean ± SEM.

**Supplementary figure 4.** (A) Gating strategy for flow cytometry analysis of IL-10^+^ monocytes in the lamina propria immune cells isolated from intestinal mucosa. (B) Percentage of IL-10^+^ cells among CD14^+^ monocytes in the lamina propria immune cells isolated from intestinal mucosa (ileum and colon) from ADK and IBD patients showed separately, stimulated for 16 h with different doses of *F. prausnitzii* EXL01 strain (Fp1, Fp10, Fp100) or LPS, as performed previously. N=9 (IBD ileum N=3; IBD colon N=2; ADK ileum N=1; ADK colon N=3). (C) Percentage of IL-10^+^ cells among CD3^+^ T and CD19^+^ B cell populations in the lamina propria immune cells isolated from intestinal mucosa (ileum and colon) from IBD and ADK patients, stimulated for 16 h, as performed previously. Data are mean ± SEM.

**Supplementary figure 5.** Over-representation analysis of the differential gene expression profiles of PBMCs between the Fp100 and Ctr conditions (A), LPS and Ctr conditions (B) and Fp100 and LPS conditions (C) at 4 h on REACTOME pathway database. Interleukin_10_Signaling pathway is encircled in green, inflammatory pathways in pink and cell metabolism-related pathways in purple.

Supplementary figure 6 . (A) Schematic representation of the two main energy metabolic pathways, glycolysis and mitochondrial respiration, including TCA cycle and oxidative phosphorylation. TCA, tricarboxylic acid cycle or Krebs cycle; ADP, adenine diphosphate; ATP, adenine triphosphate; NADH, Nicotinamide adenine dinucleotide; FADH2, Flavin adenine dinucleotide. (B) Schematic representation of the mitochondrial respiratory chain with the targets of each inhibitor used in the Seahorse Mito Stress assay. (C) Schematic representation of Oxygen Consumption Rate (OCR) evolution in time during a Mito Stress assay, showing basal respiration, ATP production, maximal respiration, spare capacity and non-mitochondrial respiration.

