## Supplementary figures and images for "*Faecalibacterium prausnitzii* induces an anti-inflammatory response and a metabolic reprogramming in human monocytes"

### Supplementary figure 1

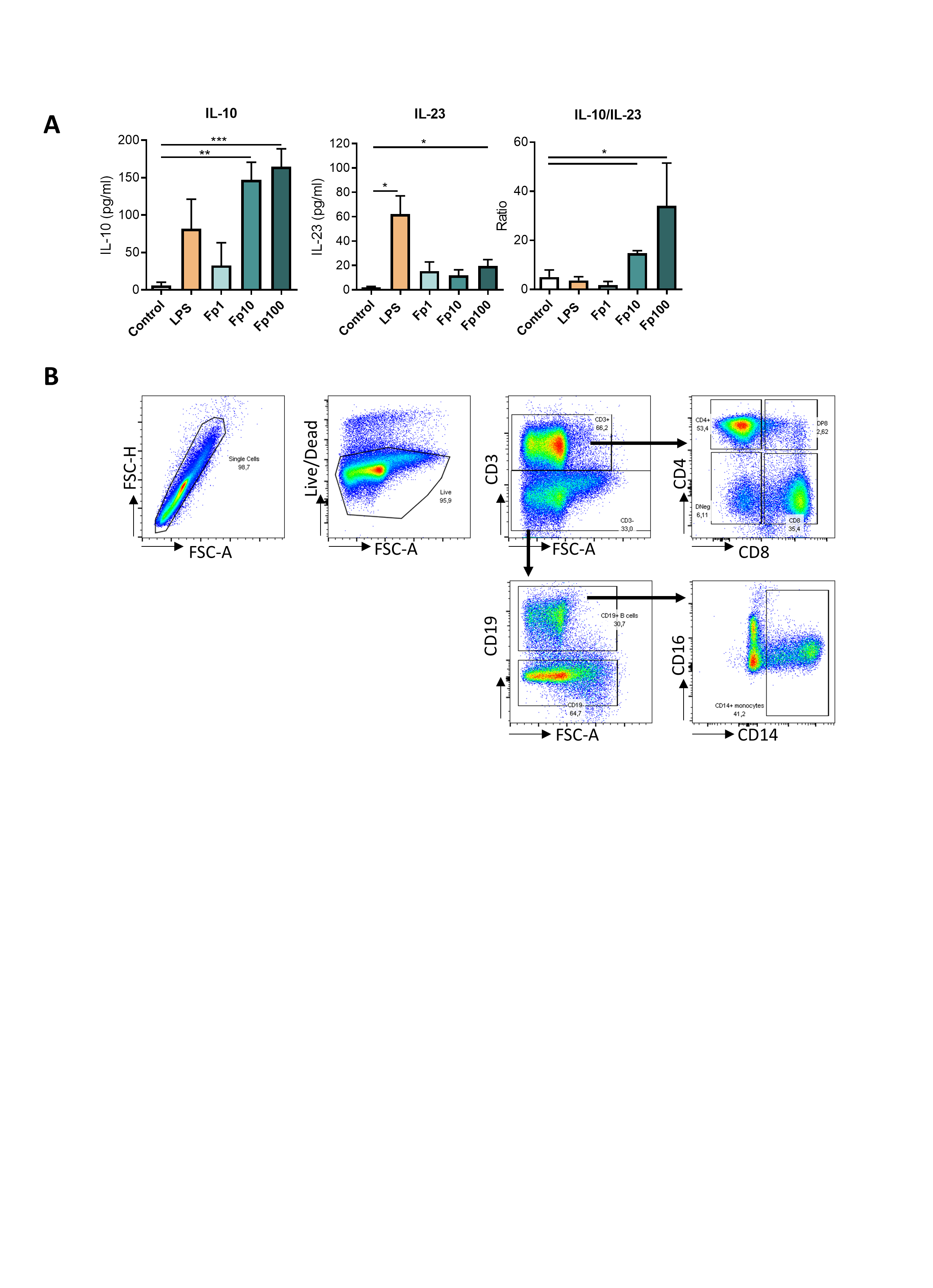

### Supplementary figure 2

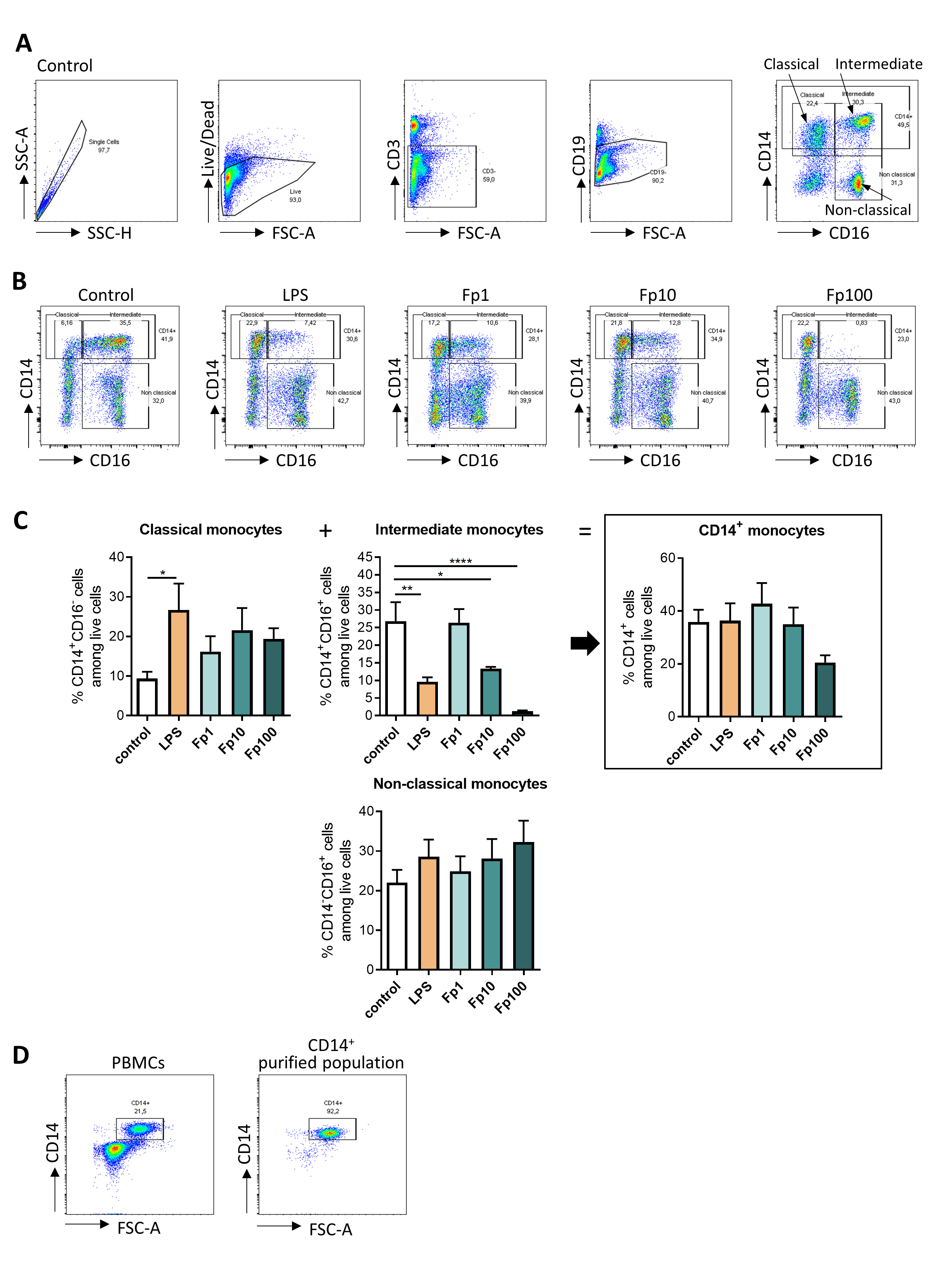

### Supplementary figure 3

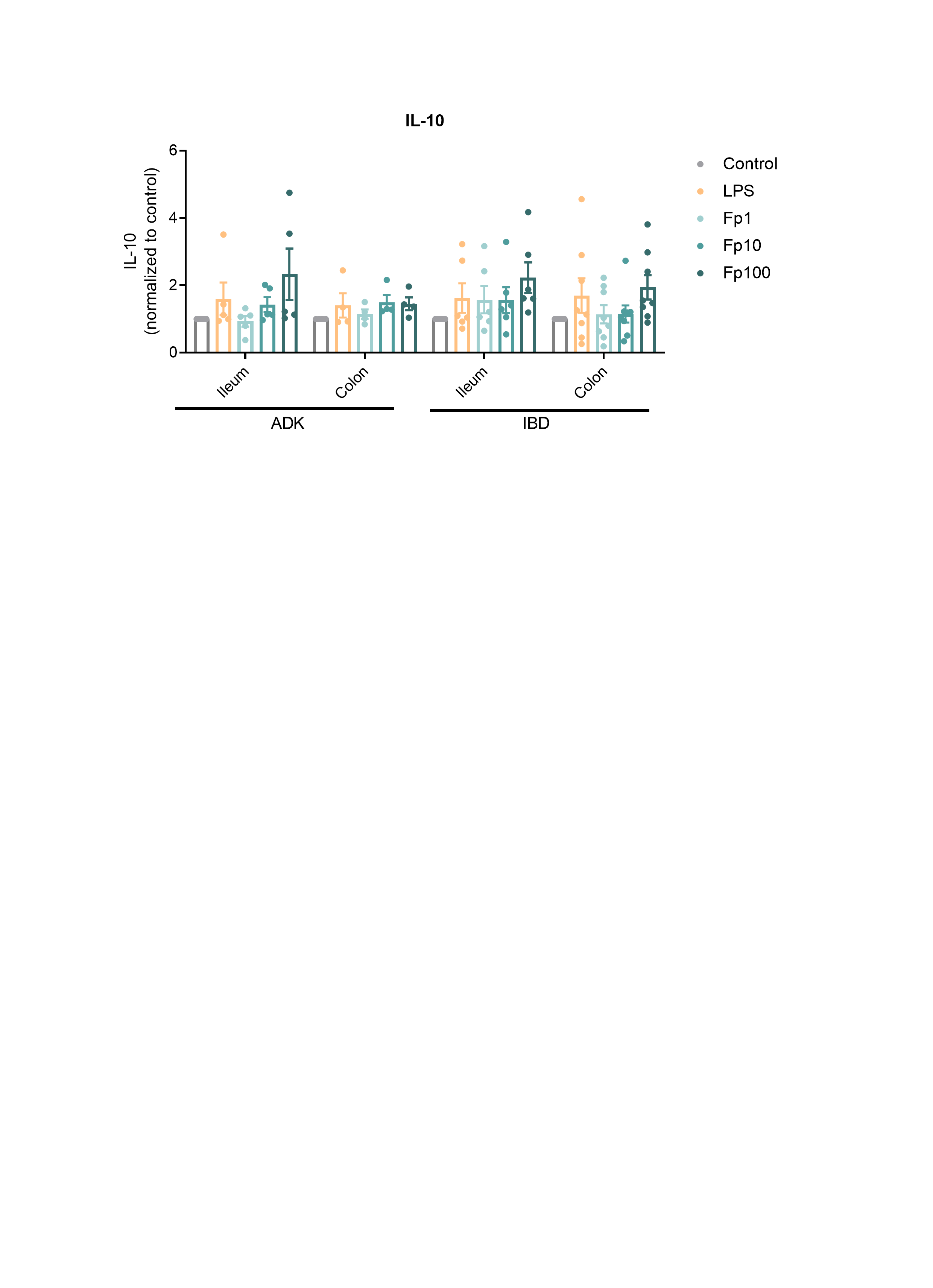

### Supplementary figure 4

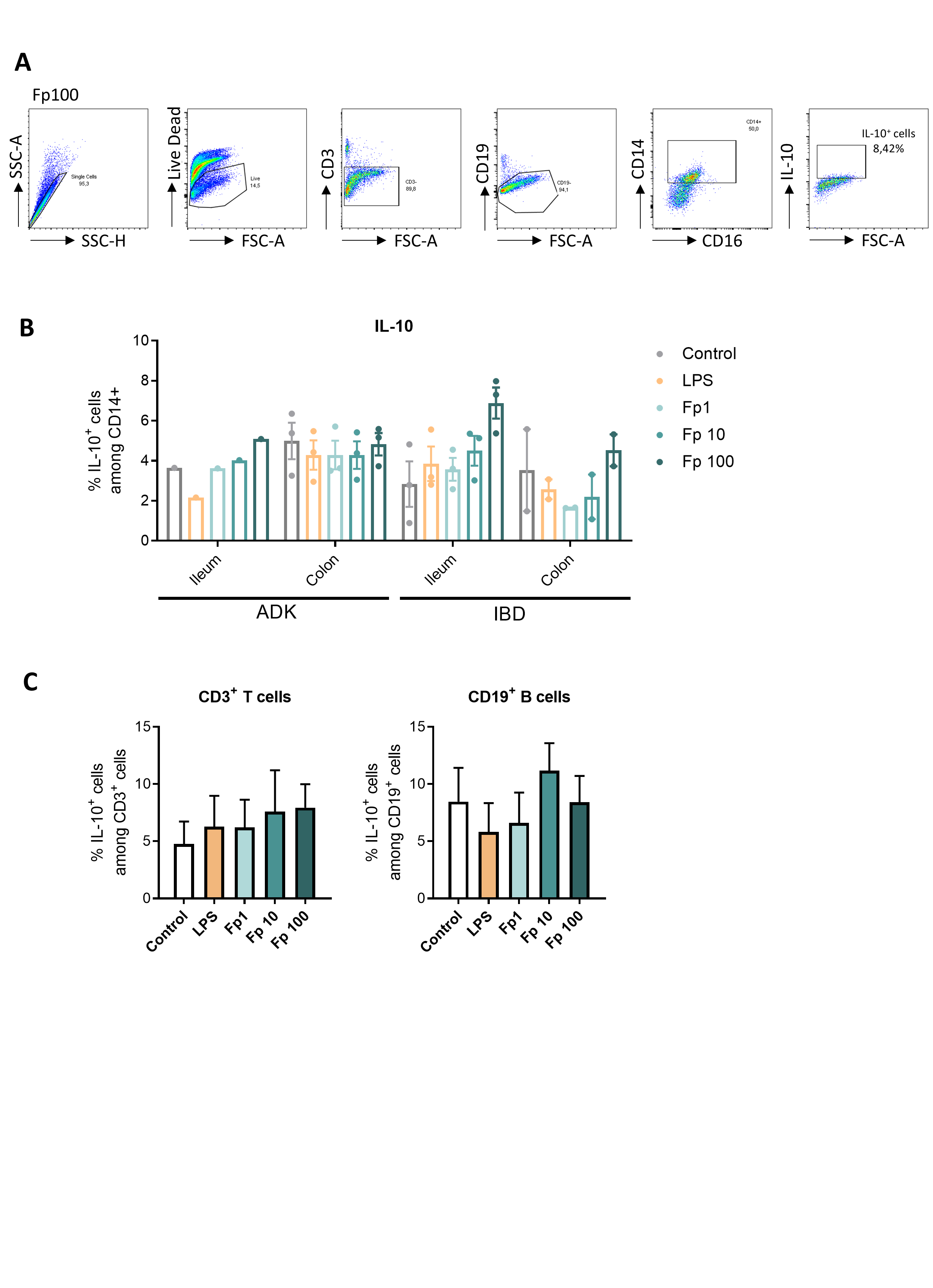

### Supplementary figure 5

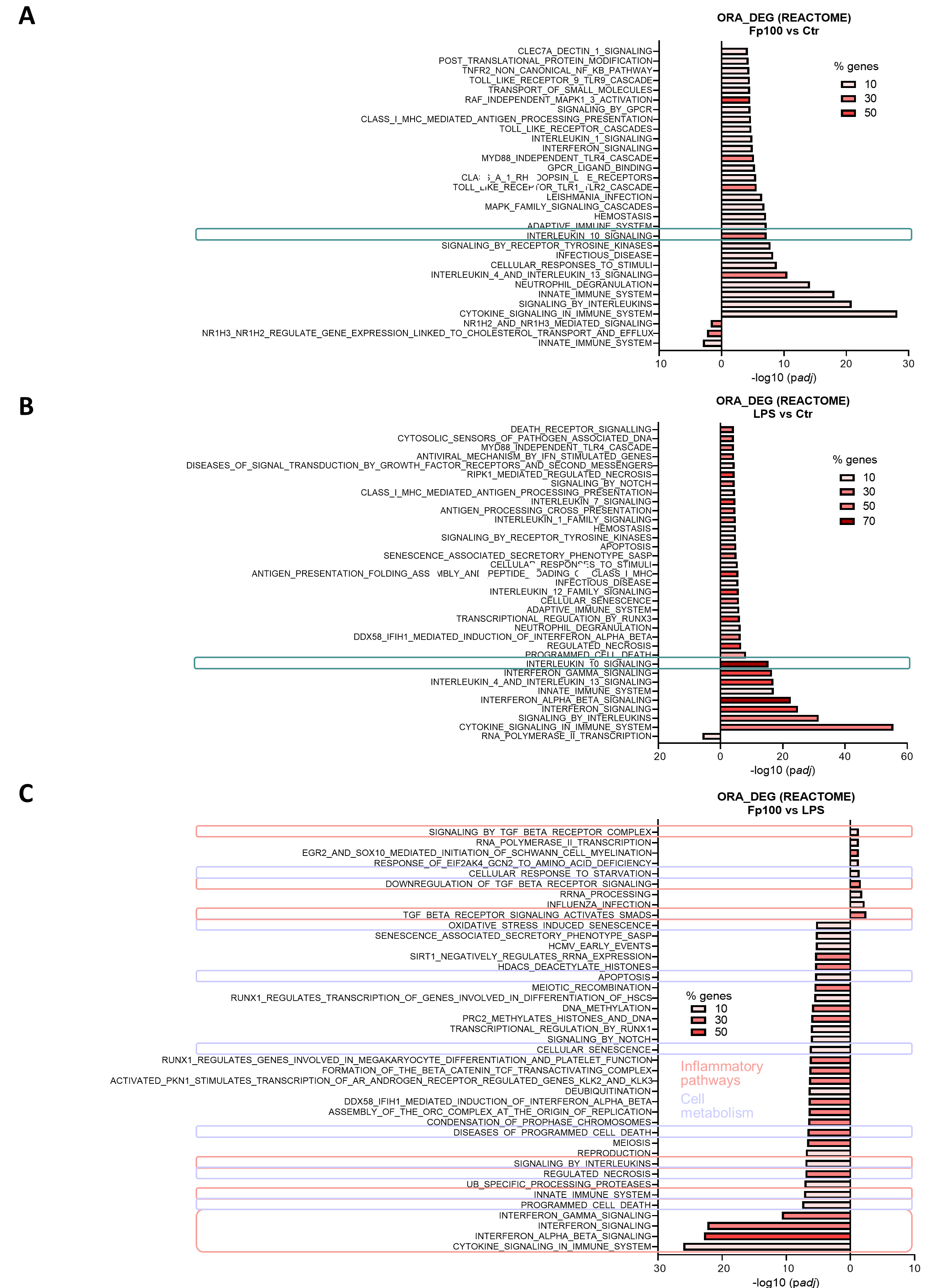

### Supplementary figure 6

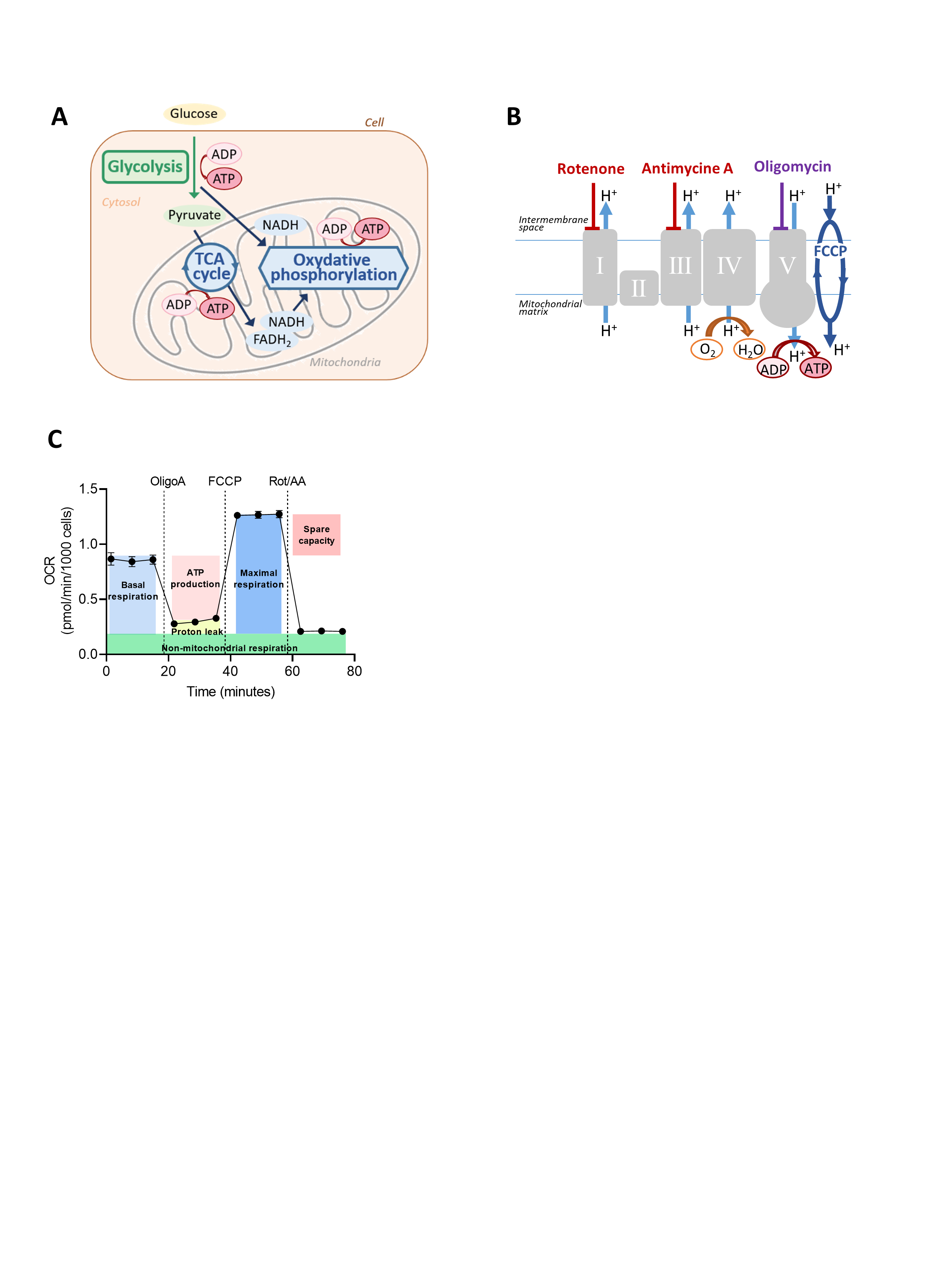
